# Identifying the diagnostic value of microRNA-421 in gastric cancer patients: a meta-analysis

**DOI:** 10.1101/468983

**Authors:** Yan Miao, Ying Zhang, Lina Wang, Lihong Yin

**Author notes:** Correspondence Author: Lihong Yin.

## Abstract

**Objectives:** Emerging evidence has shown that the expression level of microRNA-421 (miR-421) was significantly different between gastric cancer (GC) patients and healthy individuals. However, the diagnostic accuracy of miR-421 in the reports remains inconsistent. This meta-analysis aims to assess the diagnostic value of miR-421 in GC detection.

**Methods:** All related articles on miR-421 in GC diagnosis were retrieved until September 2018. The QUADAS-2 checklist was used to assess the methodological quality of each study. The diagnostic performance of miR-421 for GC were assessed by using Meta-DiSc 1.4 and STATA 14.0 statistical software.

**Results:** A total of 172 GC patients and 154 healthy controls from three articles (four studies) were enrolled in this meta-analysis. The results of pooled sensitivity, specificity, diagnostic odds ratio (DOR) with 95% confidence interval (CI) were 0.90 (95% CI: 0.85 to 0.93), 0.83 (95% CI: 0.77 to 0.87), and 37.18 (95% CI: 8.61 to 160.49), respectively. The area under the summary receiver operating characteristic curve (SROC) was 0.8977.

**Conclusions:** This study indicates that miR-421 could serve as a promising biomarker for GC detecting. Further studies are needed to verify the generalizability of these findings.

## INTRODUCTION

Gastric cancer (GC) ranks the third most common cancer in the world and the sixth leading cause of cancer-related death [1]. It appears with the highest incidence in Asia countries, particularly in China, Japan, and South Korea [2-4]. Even in Western countries, GC remains difficult to cure as most patients present in an advanced stage. Due to the invasiveness and discomfort of gastroscopy, its clinical application has been limited to a certain extent [5, 6]. Furthermore, common non-invasive markers, including carbohydrate antigen 19-9 (CA19-9) and carcinoembryonic antigen (CEA), they do not stably achieve a satisfactory sensitivity [7, 8]. Given this, it is urgent to find a novel method with better performance and minimal invasions in detecting GC.

MicroRNAs (miRNAs), which belong to a group of small (18-24 nucleotides) non-coding RNAs [9], play a pivotal role in various cell biological processes, such as proliferation, differentiation, and apoptosis [10-12]. Apart from this, increasing evidence has proved that dysregulated expression (up or down) of miRNAs usually associates with tumorigenesis [13]. Several research teams have reported that miRNAs can be stably detected in tumor tissues, cell lines, and body fluids of patients, and might be considered as potential tools for detecting related cancers [14-16]. Many studies have applied miRNAs in the diagnosis of a series of cancers, including lung cancer, breast cancer, colorectal cancer, and gastric cancer [17-20].

MicroRNA-421 (miR-421) is one of the most significantly expressed miRNAs in human cancers. The diagnostic performance of miR-421 in various cancers has been intensively investigated in the past years. Several studies indicated that miR-421 has the critical importance in GC detection [21-24]. However, the results were still inconsistent. Therefore, we aimed to collect the related studies and conduct a meta-analysis to confirm the diagnostic value of miR-421 in GC patients.

## MATERIALS & METHODS

### Search strategy

Literature retrieval was done on September 2018 through the following databases: Scopus, PubMed, Semantic Scholar, and Dimensions. The search terms were used as follows: i) terms for target tumor: “gastric cancer^∗^” or “gastric neoplasm^∗^” or “stomach cancer^∗^” or “stomach neoplasm^∗^” or “gastric carcinoma^∗^” or “stomach carcinoma” or “cancer of the stomach” or “cancer of stomach”; ii) terms for diagnostic biomarker: “microRNA-421” or “miRNA-421” or “miR-421” or “has-miR-421.” In the present study, only human studies were included. No language restriction.

### Selection criteria

The target articles should meet the following criteria: i) histopathology examination was set as the gold standard for GC diagnosis; ii) miR-421 was applied in the detection of GC patients; iii) participants were divided into two groups, case group and control group; iv) samples were collected before any treatments; v) studies had sufficient data to calculate sensitivity and specificity.

Additionally, the following items were used as the exclusion criteria: i) non-research articles; ii) duplicate articles; iii) lack qualified data; iv) participants in the study less than 30.

### Data extraction and quality assessment

Suitable data were independently extracted by two reviewers (YM and YZ) from the target articles. The following data were collected: the author’s name, publication year, number of participants, gender ratio, detection methods, and statistical results (sensitivity/specificity).

The Quality Assessment of Diagnostic Accuracy Studies 2 (QUADAS-2) checklist [25] was used to assess the methodological quality and give a corresponding score (maximum score of 7) to each study.

### Statistical analysis

The statistical analyses were performed by Meta-DiSc 1.4 and STATA 14.0 [26]. Here, the *p*-value <0.05 was considered as significant. A random effects model would be applied to calculate the following pooled values: i) sensitivity and specificity; ii) positive likelihood ratio (PLR) and negative likelihood ratio (NLR); iii) diagnostic odds ratio (DOR) when significant heterogeneity (*I^2^* index >50%) was detected [27]. The threshold effect was evaluated by Spearman correlation coefficient. If the threshold effect was absent, meta-regression would be applied to explore the possible sources contributed to heterogeneity. Furthermore, the summary receiver operating characteristic curve (SROC) was used to assess the overall diagnostic performance of the target biomarker. Deeks’ funnel plot test was used to investigate the publication bias [28].

## RESULTS

### Article selection and quality assessment

603 related articles were identified through literature retrieval, among them, 226 items were excluded as duplicates. 346 articles were excluded for various reasons (e.g., not focus on GC diagnosis) through screening the titles and abstracts. Among the left 31 articles, 27 items were excluded for lacking sufficient data through screening the full text. Finally, three articles [21-23] met the selection criteria and were included in this meta-analysis (Fig. 1).

**Figure 1.**
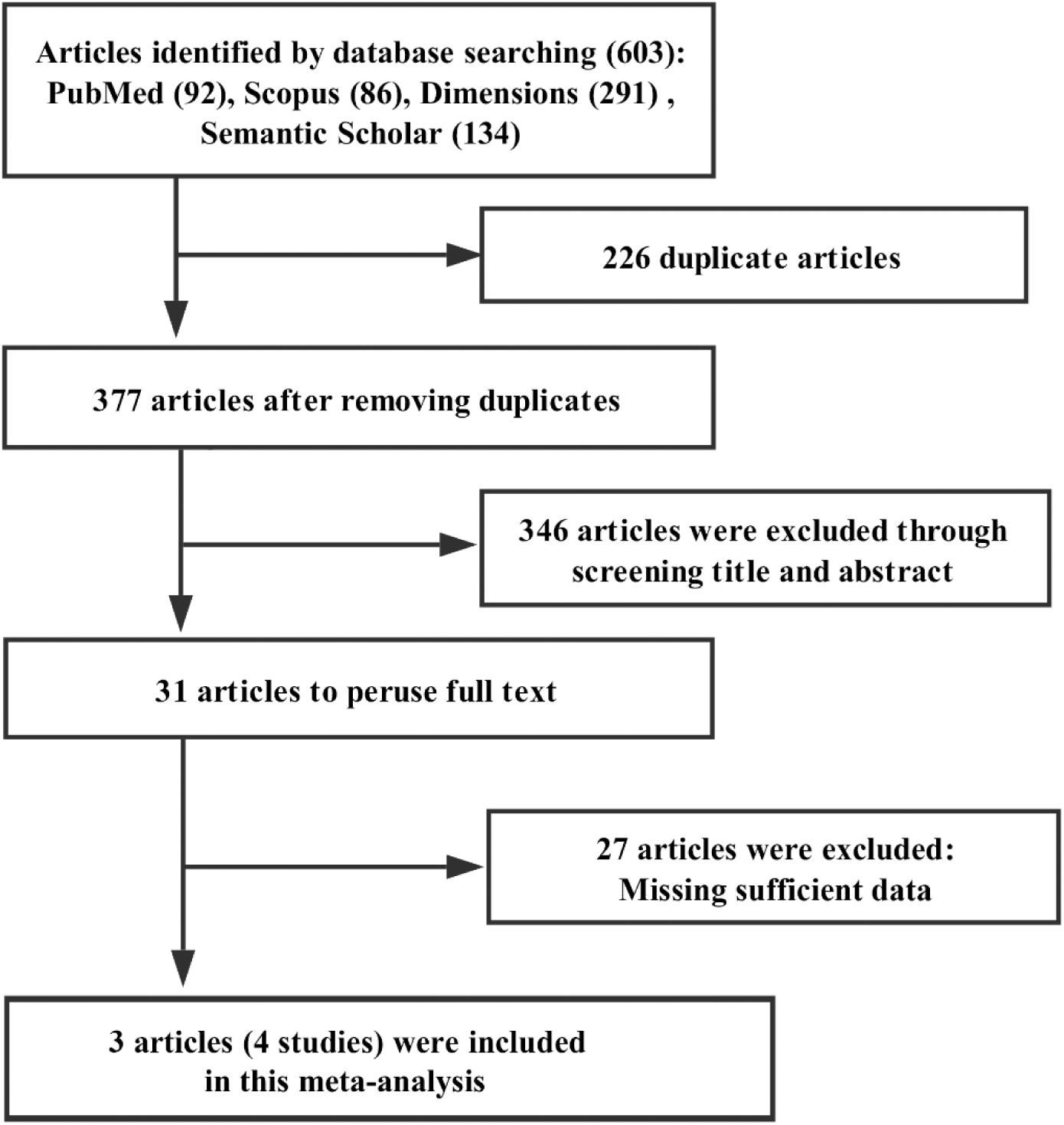
Flowchart of the article selection.

The three articles contained four eligible studies which were published between 2010 and 2014, including a total of 172 GC patients and 154 healthy controls. Histopathology examination was set as the gold standard for GC diagnosis in all of these four studies. Besides, real-time PCR (RT-PCR) was utilized to identify the expression of miR-421. Table 1 presented the detailed characteristics of these four studies, and QUADAS-2 results were listed in Table 2.

**Table 1.** Main characteristics of the studies included in the meta-analysis.

| **First author** | **Year** | **Patients (controls)** | **Stage I, II%** | **Mean age** | **Male%** | **Se%** | **Sp%** | **QUADAS-2 Score** |
| --- | --- | --- | --- | --- | --- | --- | --- | --- |
| Zhou et al | 2012 | 40(17) | 52.5 | 64.9 | 72.5 | 94.1 | 62.5 | 4 |
| Wu et al | 2014 | 90(90) | 58.8 | NR | 48.8 | 90.0 | 85.7 | 5 |
| Wu et al | 2014 | 90(90) | 58.8 | NR | 48.8 | 95.5 | 89.1 | 5 |
| Zhang et al | 2012 | 42(47) | NR | 56.8 | 65.2 | 71.4 | 71.7 | 4 |

Se, sensitivity; Sp, specificity; NR, not report

**Table 2.**
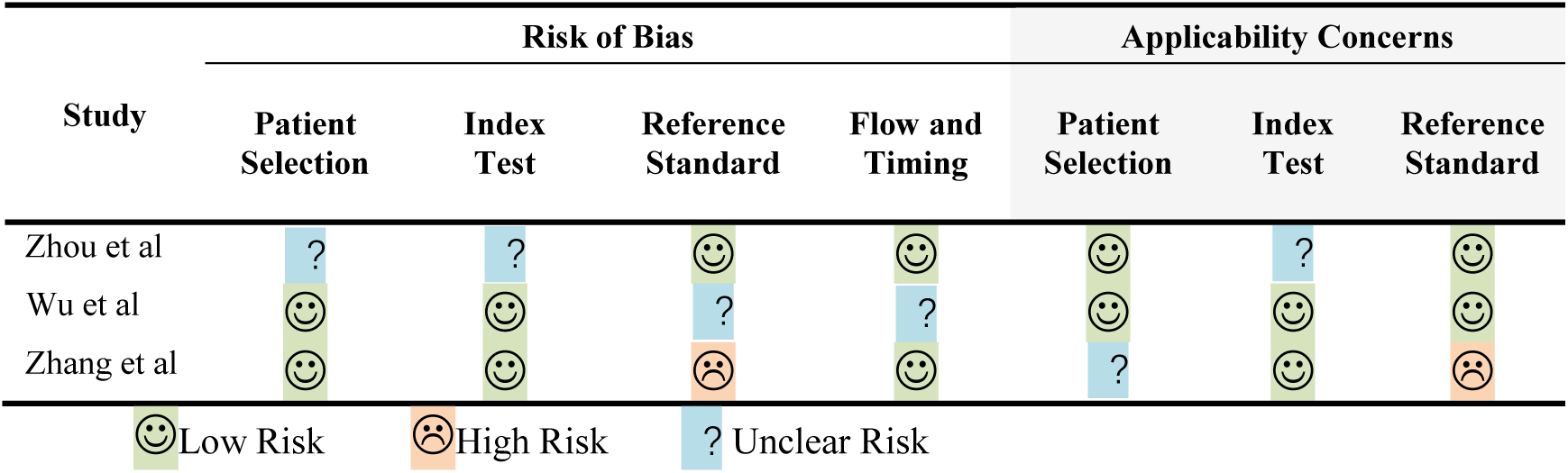
Quality assessment for the included studies.

### Data analysis

Significant heterogeneity was identified in this meta-analysis. Thus, a random effects model was applied for further analysis. The pooled sensitivity and specificity of miR-421 in GC detection was shown in Figure 2, with 0.90 (95% confidence interval (CI): 0.85 to 0.93) and 0.83 (95% CI: 0.77 to 0.87), respectively. The pooled PLR, NLR, and DOR were presented in Figure 3, with 4.40 (95% CI: 2.39 to 8.10), 0.12 (95% CI: 0.04 to 0.36), and 37.18 (95% CI: 8.61 to 160.49), separately. Furthermore, the area under the curve (AUC) of SROC was 0.8977 (Fig. 4).

**Figure 2.**
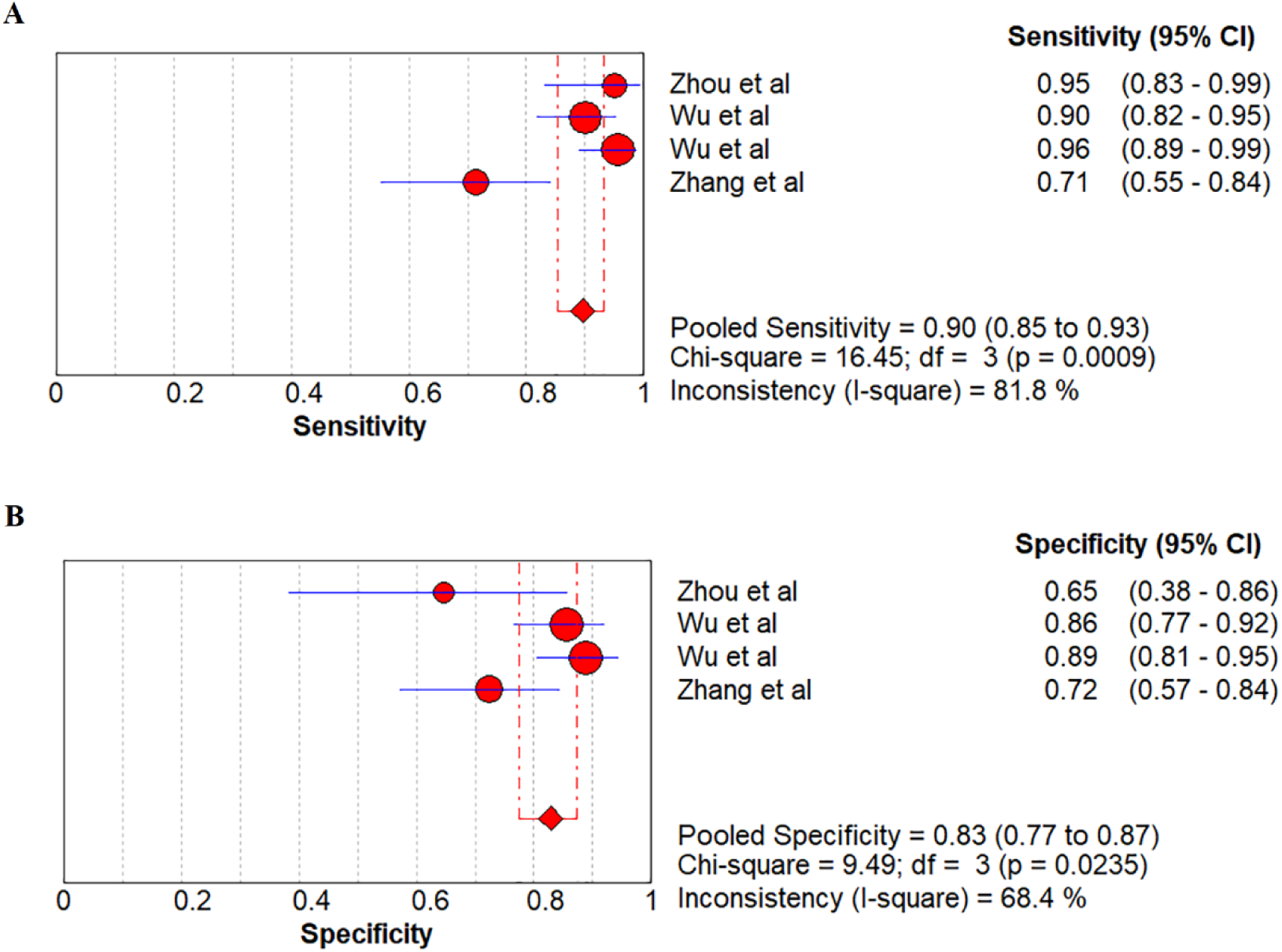
Forest plot of pooled (A) sensitivity and (B) specificity for miR-421 in detecting gastric cancer.

**Figure 3.**
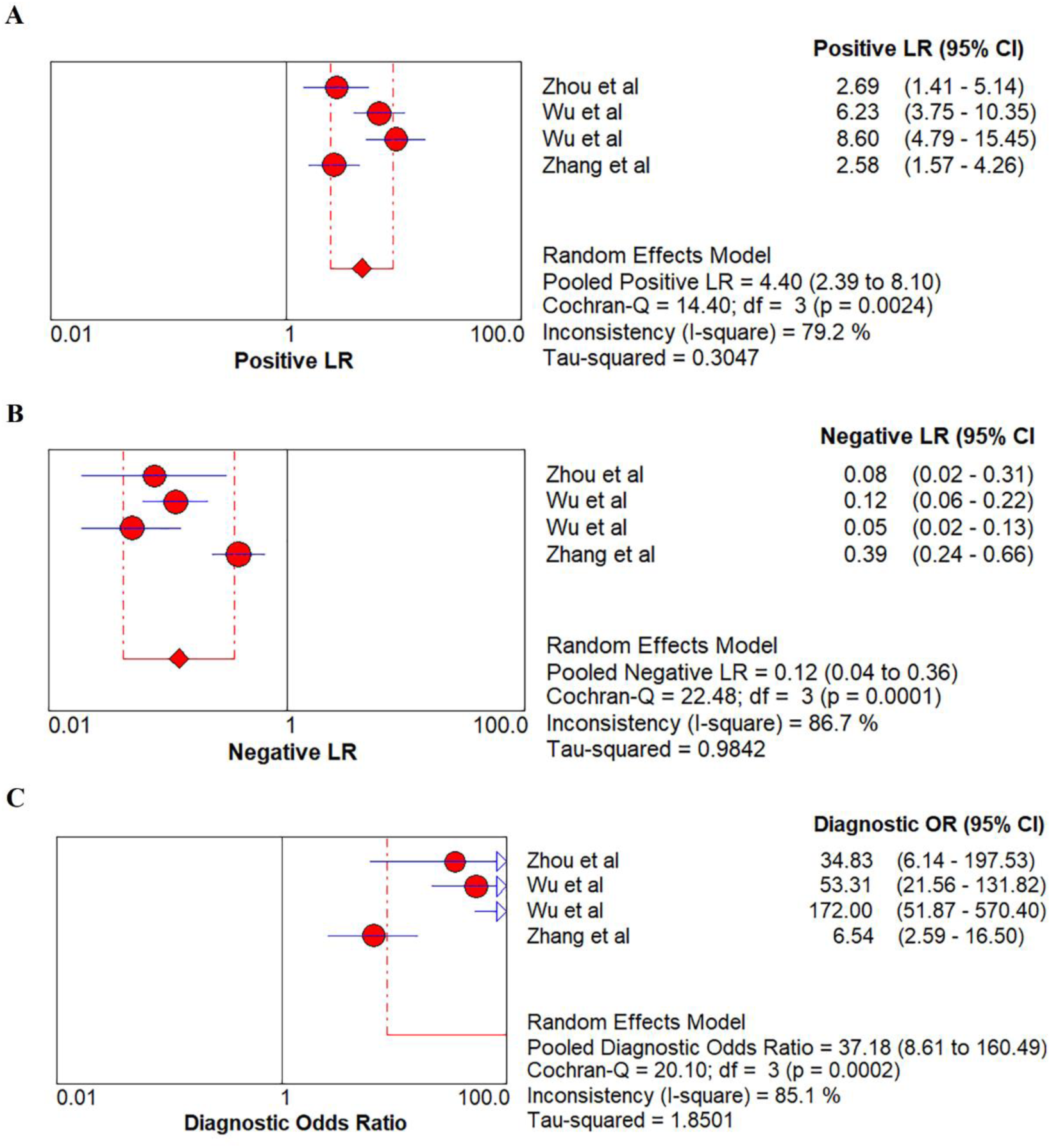
Forest plots of pooled (A) positive likelihood ratio, (B) negative likelihood ratio, and (C) diagnostic odds ratio of miR-421 in detecting gastric cancer.

**Figure 4.**
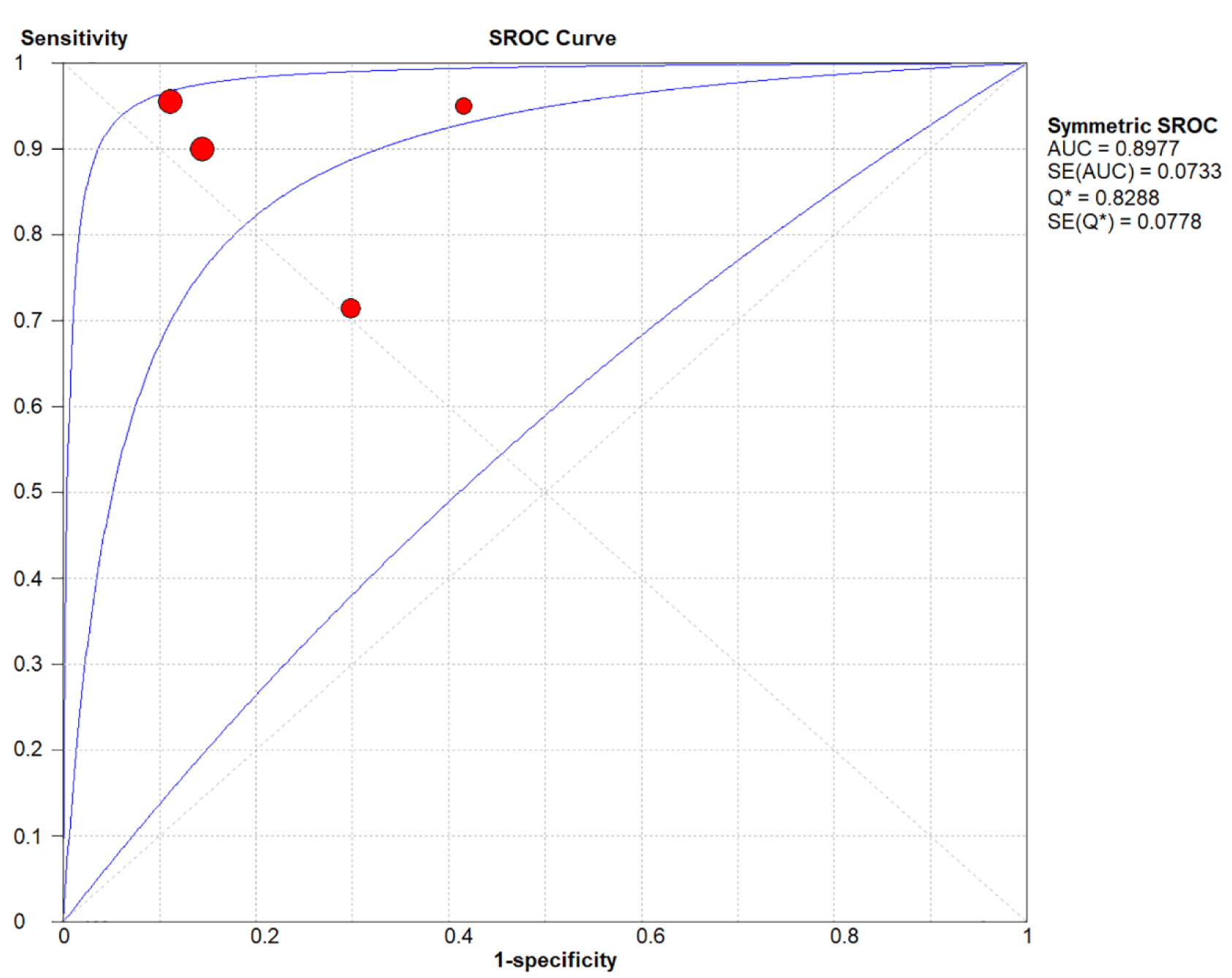
Summary receiver operating characteristic curve of miR-421 in detecting gastric cancer.

### Threshold effect and heterogeneity

The threshold effect of this meta-analysis was evaluated by the Spearman’s rank correlation analysis [26]. A correlation coefficient of −0.40 (*p*-value =0.60), besides the distribution of plane scatter did not present a typical “shoulder arm shape” (Fig. 5), suggesting that there was no threshold effect in this study.

**Figure 5.**
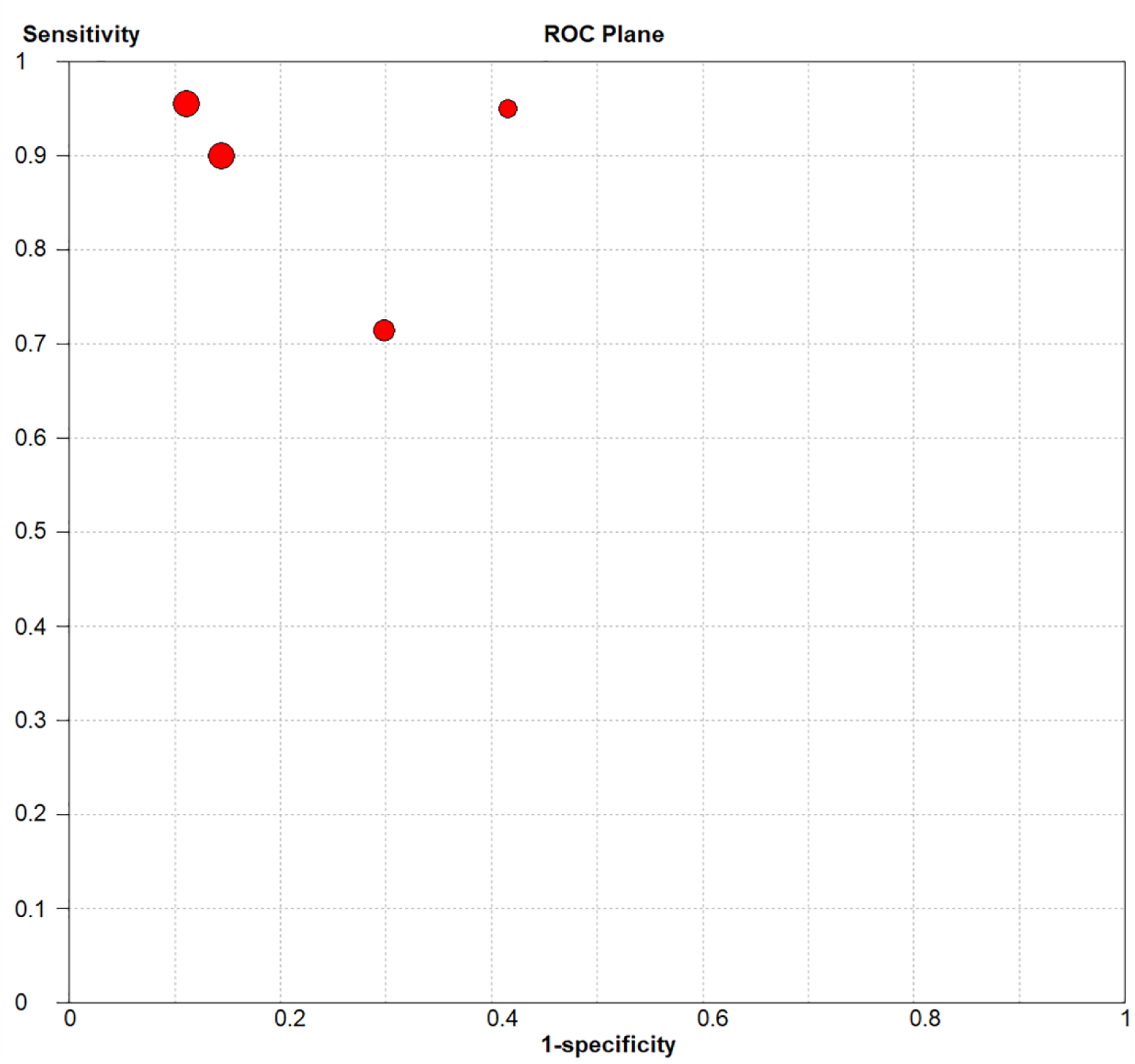
The ROC plane of the included studies.

With the absence of a threshold effect, the heterogeneity could be generated by other factors. Therefore, meta-regression analysis was applied to explore the possible factors that contributed to heterogeneity. Initially, four factors (quality, design, sample size, and gender ratio) were considered as the potential options. However, insignificant statistical differences were observed among them. The heterogeneity may be caused by other sources.

### Publication bias

To explore the impact of publication bias on included studies, Deeks’ funnel plot test was conducted in this meta-analysis. A *p*-value of 0.42 indicated that no publication bias existed in this study (Fig. 6).

**Figure 6.**
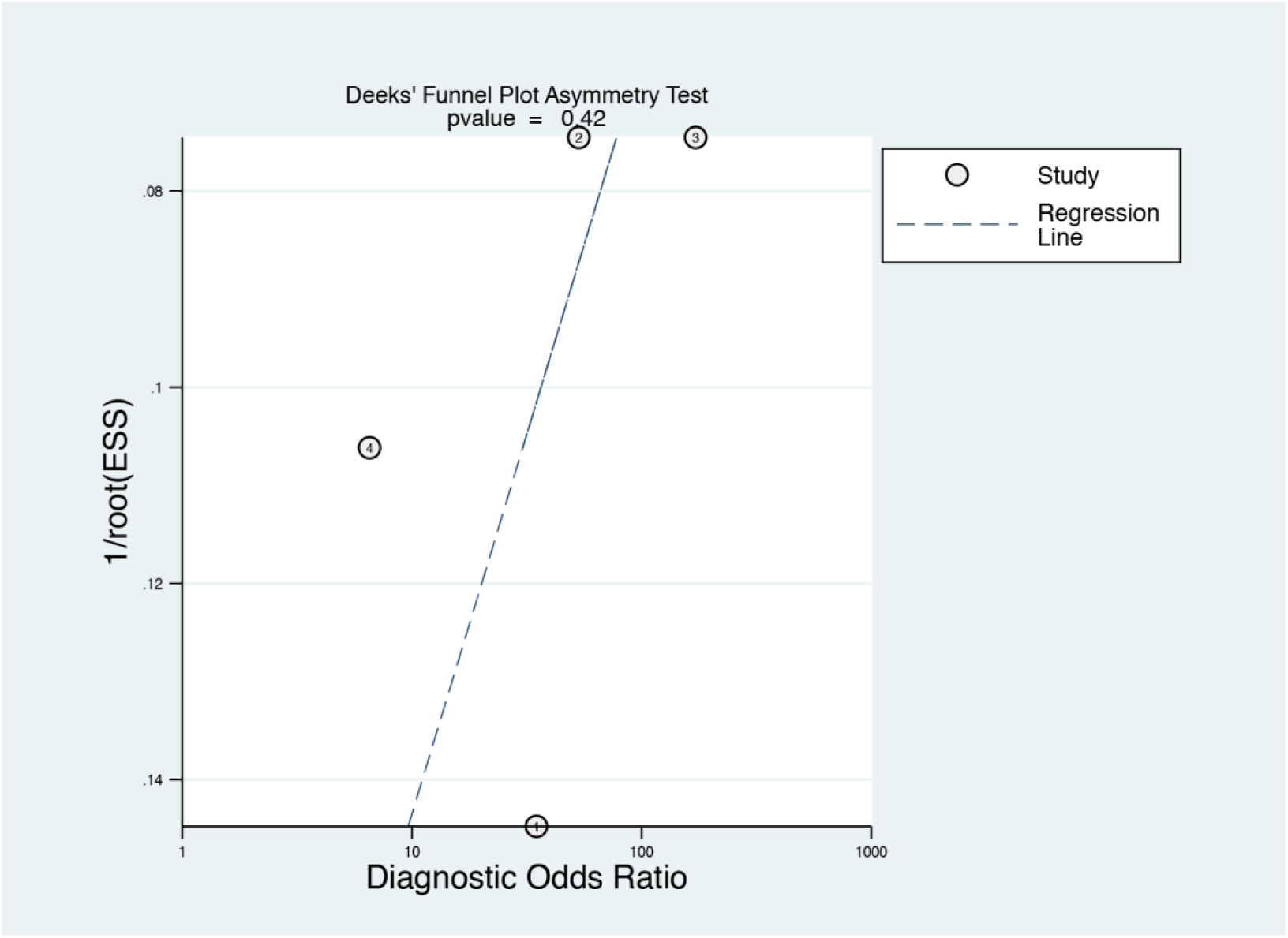
Deeks’ funnel plot test for the assessment of publication bias.

## DISCUSSION

Gastric cancer (GC) is listed as one of the most common malignancies in the world. Although the promotion of detection methods has improved the survival rate of GC patients to some extent, there are still a considerable number of GC patients diagnosed at an advanced stage with a poor prognosis [29]. Early diagnosis is not only conducive to the treatment of disease but also helps to improve the later rehabilitation [30]. There is a need to find a novel and non-invasive method to improve GC detection. During the past years, several studies reported that miR-421 has the diagnostic value in detecting GC. Therefore, this study was dedicated to verifying the diagnostic accuracy of miR-421 in patients with GC.

In the present study, the overall sensitivity and specificity of miR-421 in GC detection were 0.90 (95% CI: 0.85 to 0.93) and 0.83 (95% CI: 0.77 to 0.87), respectively, demonstrating that miR-421 had the potential to be a diagnostic biomarker for GC patients. The DOR that combines the advantages in both sensitivity and specificity is an overall indicator of diagnosis performance. If the value of DOR is greater than 1, the target biomarker would have a better performance in discriminating patient cases and healthy controls. On the contrary, if the value of DOR is less than 1, the likelihood of healthy individuals being misdiagnosed would be increased. Here, the overall DOR of miR-421 was 37.18 (95% CI: 8.61 to 160.49), suggesting that miR-421 had an ideal effect on detecting GC. Regarding the AUC, the closer a value of AUC is to 1, the better the diagnosis efficiency will be. Generally, the value of AUC can be divided into three grades: i) value is greater than 0.9 (excellent efficiency); ii) value is between 0.7 and 0.9 (moderate efficiency); iii) value is between 0.5 and 0.7 (mild efficiency). In this meta-analysis, an AUC of 0.8977 implied a good diagnosis efficiency. Thus, miR-421 could be a promising diagnostic biomarker for GC patients.

Regarding miR-421 itself, it has some advantages as a novel biomarker in GC diagnosis. miR-421, which belongs to the miR-374b/421 cluster [31], is stably expressed in either peripheral blood or tissues. Compared with the standard histopathology examinations, miR-421 has minimal invasion and better convenience. For GC patients, gastroscopy is a routine diagnostic method. However, many patients are afraid or refuse to choose it as a means of early diagnosis due to its complicated procedures and discomfort experience, which limits its widespread applications in the clinic. Fortunately, miR-421 shows a better diagnostic performance (both sensitivity and specificity were above 0.8) than some traditional biomarkers, including CEA and CA19-9 [32, 33]. It is expected to be further applied in related diagnosis.

There were several limitations existed in our study. Firstly, the differential expression of miR-421 has been previously reported in other cancer, including breast cancer, cervical cancer, and hepatocellular carcinoma [34-36]. The results demonstrated that miR-421 might not be particularly relevant to GC itself, it is also associated with the progression of other common cancers. miR-421 is still a promising biomarker for GC detection, but it can combine with other biomarkers to extend its clinical applications. Secondly, the emergence of miR-421 has indeed caught the attention of researchers, but limited studies have focused on this topic. Based on this, only a few number of articles were included in this meta-analysis, which caused a relatively small sample size. We could not achieve a subgroup analysis to explore the other sources of heterogeneity, either. Thirdly, GC is a common cancer in Asia regions, especially in China [37]. Coincidentally, the appropriate studies included in this meta-analysis were all from China. Therefore, further multi-center research are needed to verify our findings, especially the use of miR-421 in detecting GC in different racial patients. Finally, these four studies only recruited healthy individuals as controls, which may cause the overestimation of the sensitivity.

## CONCLUSIONS

In summary, this meta-analysis evaluated the potential role of miR-421 in GC patients. The findings indicated that miR-421 showed a good diagnostic accuracy and could regard as a promising biomarker in GC detection. Furthermore, large-scale and well-designed studies are needed to conduct to confirm the wide-ranging use of miR-421 in clinical practice.

## ACKNOWLEDGMENTS

We thank Ms. Lina Wang for her help in data analysis.

